# Early-life gut microbiome and stress-axis perturbations dysregulate systemic, mucosal, and brain immunity

**DOI:** 10.1101/2025.03.19.644259

**Authors:** Van A. Ortega, Michelle R. Asbury, Colin Mackenzie, Fernando A. Vicentini, Bilal Hafeez, Haonan Duan, Emily M. Mercer, Erik van Tilburg Bernardes, Kristen Kalbfleisch, Jumana Samara, Veronika Kuchařová Pettersen, Daniel Figeys, Keith A. Sharkey, Marie-Claire Arrieta

## Abstract

**Background:** Early-life disruptions to the gut microbiome and stress-axis significantly influence the development of immune, neuroendocrine, and other physiological systems. However, the precise microbial species and pathways mediating these effects remain poorly characterized. Using a murine model, we investigated the individual and combined effects of early-life antibiotic exposure and chronic stress on gut microbiota composition, short-chain fatty acid (SCFA) production, hypothalamic-pituitary-adrenal (HPA) axis activity, and systemic, mucosal, and neuroimmune responses.

**Results:** Broad-spectrum antibiotic treatments severely reduced microbial diversity and SCFA concentrations, with changes persisting into adulthood. Chronic early-life stress exerted more modest but notable effects, reducing key SCFA-producing taxa and impacting microbiome metabolic output. Combined disruptions led to altered microglial active phenotype and cytokine profiles, impaired immune cell populations, and suppressed HPA axis activity. Multi**-**omic correlational analyses revealed strong associations between SCFAs, specific gut microbes, and immune responses, implicating SCFAs as critical mediators of gut-brain communication. Notably, antibiotic exposure exacerbated susceptibility to allergic airway inflammation, highlighting the systemic consequences of early-life microbiome disturbances.

**Conclusions:** These findings demonstrate that early microbial perturbations impair neuroimmune maturation, HPA axis regulation, and host resilience to inflammatory diseases. Our study underscores the importance of preserving the early-life microbiome to support long-term immune and neurodevelopmental health, offering insights into potential therapeutic interventions for mitigating the impact of early-life microbiota disruptions.

## Introduction

The gut microbiome is now recognized as an essential exposure in early-life host development, with microbial colonization during infancy not only coinciding with, but also influencing key developmental windows for brain neural networks and host immunity. These time-delimited microbial colonization patterns are pivotal in regulating the maturation and functionality of these systems through communication pathways in the gut-brain axis.^1,2^ This axis functions bidirectionally, as evidenced in animal models where exposure to stress shifts the compositional profile of gut microbes,^3^ even in fear-evoked rodents that only witnessed stress.^4^

Conversely, neonatal germ-free (GF) mice (i.e., born sterile and lacking a microbiome during development) display altered activity of their hypothalamic pituitary adrenal (HPA) stress axis when stimulated, but which can be rescued when colonized with *Bifidobacterium* spp. during early life, demonstrating the time-sensitive nature of microbial regulation during this developmental window.^5^ In humans, antibiotics are often administered to hospitalized infants to help combat nosocomial infections that their underdeveloped immune systems cannot yet manage. Hospital-acquired infections are especially problematic for preterm infants who are not only at greater risk of neuroimpairment and inflammatory disorders owing to their early birth and stressful hospital experiences,^6^ but the microbiome-altering effects of antibiotics themselves have also been associated with the development of inflammatory disorders like asthma,^7^ even in children without structural or functional lung damage,^8^ suggesting that altered microbial colonization in early-life may also contribute to maldevelopment. Thus, a complex interplay exists between proper establishment of the postnatal gut microbiome and early-life exposures to stress and other factors (e.g. postnatal antibiotics) that dictate long-term neuronal and immune health.

In vertebrates, the HPA stress axis regulates the complex, but highly coordinated release of autonomic and neuroendocrine catecholamines and glucocorticoid hormones that provoke multifaceted physiological changes aimed at restoring homeostasis. The extensive impact of HPA-axis dysregulation on human disease ranges from neurological and psychiatric disorders like anxiety, depression, and multiple sclerosis, to systemic inflammatory diseases such as allergic asthma.^9^ In mice, for example, exposure to stress during early-life increases asthma severity,^10^ while early-life treatment of stressed animals with probiotics ameliorates lung inflammation.^5^

The biological connections that mediate crosstalk between the gut microbiome and the brain involve neural and immune pathways, with microglial cells playing a critical role as a biological bridge in microbiome-mediated stress activation.^11^ Overactive microglia can contribute to stress-related anxiety and depressive disorders,^12,13^ as well as cognitive deficits and neuroinflammation following septic infection as adults^14^ and during perinatal periods.^15^ Chemically blocking or eliminating stress-experienced microglia prevents both immune reactivity in the brain and blocks anxiety recurrence following a stress challenge.^13^ Microglia are also receptive to peripheral and gut-derived chemical messengers such as cytokines, chemokines, and other neuroactive molecules like serotonin, melatonin, and gamma-aminobutyric acid (GABA), including those produced in the gut.^16−19^ Microglia from GF mice have chronically activated and immature phenotypes^20^ with globally defective immune responses,^16^ suggesting that disrupted neonatal microglia may contribute to GF mice having altered synaptic networking, poorer sociability, memory, and decreased anxiety-like behaviours.^21^

Despite the growing evidence linking the microbiome to immune and neurodevelopmental responses, the microbial species and pathways involved in this connection remain underexplored. Here, we examined the individual and combined effects of early-life chronic stress and antibiotic treatments on microbiome community, its production of key metabolites during developmental periods (21 days postnatally) and adulthood. We further investigated the outcomes of these microbiome disruptions on host microglia activity, HPA axis function, as well as systemic, intestinal and lung immunity.

## Methods and Materials

### Mouse husbandry

Six-week-old male and female C57BL/6 mice were purchased from Charles River (Canada) and housed in the Clara Christie Centre for Mouse Genomics at the University of Calgary. Mice were kept under 12-h light/12-h dark cycles with a maximum of 5 animals per cage and with ad libitum access to water and sterile mouse chow. All mouse husbandry procedures were completed under the approval of the Animal Care Committee protocol AC19-0112. C57Bl/6 mice pups were bred in-house before being grouped into six treatment groups.

### Experimental design

Six-week-old mice were paired in cages for the purpose of breeding the pups that would be grouped into the different treatments. Pregnant dams were given standard food (LabDiet, Pico-Vac Mouse Diet 20 5062, CA, USA) and water until the birth of their pups. Pups were separated into three antibiotic treatment groups: water (as a control), Augmentin, and a broad-spectrum antibiotic cocktail to expose them to different degrees of microbial alterations during early life.

Pups were treated with antibiotics via nursing on treated dams starting from birth until postnatal day 21 (P21). Water-treated dams were given sterile water, while Augmentin-treated dams were given amoxicillin/clavulanate (0.2 mg/mL) (MiliporeSigma, Burlington, MA, USA)^22^, a commonly prescribed neonatal and infant antibiotic used to treat various respiratory tract and nosocomial-related anerobic bacterial infections.^23,24^ The dams treated with the broad-spectrum antibiotic cocktail were given a solution of vancomycin (0.25 mg/mL; Cayman Chemical, Ann Arbor, MI, USA), gentamicin (0.5 mg/mL; MiliporeSigma), metronidazole (0.5 mg/mL; Cayman Chemical), neomycin (0.5 mg/mL; Cayman Chemical), and ampicillin (0.5 mg/mL; Cayman Chemical)^22^, a commonly used combination of antibiotics for treating various infections.^24^ Both antibiotic treatments were dissolved in sterile water beginning at birth until P21. All water treatment bottles were changed every three days.

Pups were further assigned to either a stressed or unstressed group, with the stressed group undergoing daily maternal separation from P7 until P21, while the unstressed group was not separated from their mothers and remained under standard rearing conditions. For the stressed group, mothers were removed daily from their cages and placed alone into separate cages, while their pups remained in their original cages on heating mats set to 37°C during the separation period. The maternal separation also involved increasing separation times starting from 15 min on P7, 20 min on P8, 30 min on P9, 1 h on P10, and 3 h per day from P11-P21, at which time pups are normally weaned from their mothers and become independent adolescent feeders. The increasing separation times reflect our attempts to lessen maternal cannibalization of young pups, which can occur when dams are overly stressed. At the end of each separation, mothers were placed back into their original cage with their pups.

### Tissue collection

On day P21, half of the litter for each treatment group (N=8-10/group) were euthanized in order to collect tissues for analysis. Fecal pellets were collected before the mice were anesthetized and a blood sample was collected through cardiac puncture while under isoflurane anesthesia (1L/min of O_2_ with 5% isoflurane gas for several min until righting and touch reflex were absent and breathing rate was slowed), followed by cervical dislocation. Fecal and blood samples were kept on ice until long-term storage at -80°C for microbiome and corticosterone analysis, respectively. Body hair and whiskers were collected using scissors and electric hair clippers and stored in the dark in paper envelopes for corticosterone analysis. The spleen was removed and stored in incomplete Roswell Park Memorial Institute (RPMI) 1640 GlutaMAX cell culture media (Gibco^TM^, Thermo Fisher Scientific, Waltham, MA, USA) on ice for splenocyte culture. Samples of saline-flushed colon (approx. 8 - 10 cm) from the large intestine were collected and flash frozen on dry ice before being stored long-term at -80°C to determine inflammatory cytokines. The brain was then removed and stored in 2mL of 1x Hank’s Balanced Salt solution (HBSS) without Ca^2+^/Mg^2+^ on ice for isolation of microglial cells. The remaining animals continued in the experiment undisturbed under standard rearing conditions and provided with sterile water and rodent chow until behaviourally assessed at P60, and then euthanized at P80 following induction of allergic lung inflammation (see below). Lung tissue, bronchoalveolar lavage fluid (BALF) for lung inflammation, plasma and hair for corticosterone measurements, as well as stool pellets for microbiome and metabolomic analysis were collected at this timepoint to measure adult tissue responses to early-life treatments.

### Fecal DNA isolation and qPCR analysis

DNA was extracted from fecal samples using the Qiagen DNeasy Powersoil Pro Kit (Qiagen, Germany) according to the manufacturer’s instruction with the following modifications. The fecal sample was lysed using a TissueLyser (Qiagen) for 30 seconds at 30 Hz, and repeated after changing the orientation of the samples. DNA was eluted from the column by adding a total elution volume of 35 μL of polymerase chain reaction (PCR) grade water to the spin column, then spun at 15,000 g for 1 minute (21°C). DNA concentrations were determined using a NanoDrop Lite Spectrophotometer (Thermo Fisher Scientific, Waltham, MA, USA), with PCR grade water used as blank measurements.

DNA concentrations were determined by quantitative amplification of bacterial 16S rRNA genes, as previously described for qPCR.^25^ For qPCR analysis, 10 µL reactions consisted of 5 µL of iQ SYBR Green Supermix (BioRad Laboratories, Hercules, CA, USA), 0.5 µL of each primer (universal 16S primers [U16Sfwd: 5’-TCC TAC GGG AGG CAG CAG T-3’ and U16Srvs: 5’- GGA CTA CCA GGG TAT CTA ATC CTG TT-3’])^25^ at 10 μM, 2 μL nuclease-free water, and 2 μL of 1 ng/μL dilution of extracted template DNA. Thermocycler conditions, performed on the StepOne Plus Real-Time PCR System (Applied Biosystems, Foster City, CA, USA), consisted of an initial 5 min heating step at 94°C, followed by 40 cycles of 94°C for 15s, 60°C for 30s, and 72°C for 30s, and a final melt curve consisting of a single cycle of 95°C for 15s, 60°C for 1 min, 95°C for 15s, and 60°C for 15s. All samples were run in triplicate and concentration were calculated based on standard curves generated using bacterial genomic DNA extracted with Qiagen DNeasy PowerSoil Pro kit as described above.

### 16S rRNA gene sequencing

Fecal microbial DNA was further used to amplify the V4 region of the bacterial 16S rRNA gene to determine microbial composition. Diluted DNA was sequenced at Microbiome Insights (University of British Columbia, Vancouver, Canada) where 16S rRNA gene amplicon libraries were prepared and amplified for sequencing using Phusion Hot Start II DNA Polymerase (Thermo Fisher Scientific). PCR reactions were cleaned up, normalized using the high-throughput SequalPrep^TM^ Normalization Plate Kit (Thermo Fisher Scientific), and quantified with the KAPA qPCR Library Quantification kit (Roche Sequencing Solutions, Pleasanton, CA, USA). Controls without template DNA, and mock communities with known proportions of selected bacteria within the microbial community, were included in the PCR and downstream sequencing steps to control for microbial contamination and to verify the bioinformatic analysis pipeline. The pooled and indexed libraries were denatured, diluted, and sequenced in paired-end mode on an Illumina MiSeq (Illumina Inc., San Diego, CA, USA).

### Short-chain fatty acid (SCFA) determination in stool

Mouse stool pellets were weighed, and 50% aqueous acetonitrile was added to each sample depending on sample weight to have sufficient supernatant for measurement purposes. The samples were then homogenized using mill homogenizers (MP Biomedicals, Santa Ana, CA, USA) and subsequently centrifuged at 16,000 x g and 10°C for 10 min. The supernatant was utilized for the measurement of SCFAs. The levels of SCFAs were quantified using a derivatization method, as published in a previous study.^26^ Briefly, the supernatant (40 μL) or SCFAs standard solution was reacted with 20 μL of 200 mM 3-nitrophenylhydrazine (3NPH) and 20 μL of 120 mM of N-(3-dimethylaminopropyl)-N′-ethylcarbodiimide hydrochloride-6% pyridine solution at 40 °C for 30 min. The samples were diluted by 10% aqueous acetonitrile and mixed with ^13^C_6_-3NPH-labeled SCFAs as internal standards. The mixtures were analyzed on an Orbitrap Exploris 480 mass spectrometer (Thermo Fisher Scientific) in PRM mode equipped with a tip column (75 μm inner diameter ×15 cm) packed with reverse phase beads (3 μm/120 Å ReproSil-Pur C18 resin, Dr. Maisch HPLC GmbH, Ammerbuch-Entringen, Germany). The concentrations of SCFAs were calculated based on the standard calibration curve using internal standards.

### Corticosterone determination in plasma and hair

Corticosterone analysis was performed on P21 and P80 plasma and hair (body hair and whiskers) samples using the Oxford Biomedical Research Corticosterone enzyme-linked immunosorbent assay (ELISA) kit (Cat#: EA66, Oxford Biomedical Research, MI, USA). For plasma samples, 50 µL of the ethyl ether-extracted corticosterone from each sample was added to a 96-well plate in duplicate and the ELISA was run according to the manufacturer’s instructions. Briefly, diluted corticosterone-HRP conjugate was added to wells containing sample or standard, and following a 1 h incubation, 3,3′,5,5′-Tetramethylbenzidine (TMB) substrate was added and further incubated to generate an in-well colorimetric reaction that was measured on a spectrophotometer plate reader at 650 nm wavelength. To prepare corticosterone samples from hair, we followed protocols from previously published reports^27^ with some modifications. These included repeated wash steps in High Resolution Gas Chromatography (HRGC) methanol to clean hair of contaminants, grinding hair in a ball mill homogenizer (10 min at 30 Hz for 25 mg sample) to powderize it, further cleaning in methanol, and overnight rotator incubation followed by centrifugation (2150 x g; 15 min at RT) to collect supernatant. The methanol-suspended hormone was then repeatedly concentrated and dried in glass test tubes using an N-Evap system compatible with a nitrogen tank. The extracted hormone was reconstituted in a small volume (0.2 mL) of extraction buffer (overnight at 4°C), as part of the Oxford Biomedical ELISA assay to measure accumulated corticosterone levels in hair.

### Microglia isolation

Microglia were isolated from the whole brain of animals based on the protocol outlined in Andonegui et al.^14^ with modifications. Whole brains were homogenized and digested while rotating for 30 min at 37°C in a digestion solution (1x HBSS without Ca^2+^/Mg^2+^, 0.001% of DNase 1, and 0.05% of collagenase). Following the incubation, 1x HBSS (7 mL, 21°C) was added to the solution and pelleted after spinning for 10 min (1800 rpm, 4°C). The supernatant was discarded, and the pellet was resuspended in 4 mL of percoll (70%). The 70% percoll was overlaid with 4 mL of 30% percoll, then another layer of 37% percoll. The gradient was then spun for 20 min (1800 rpm, 21°C, no break). The interface layer between the 70% and 30% percoll contained the microglial cells and was transferred to a 3 mL fluorescence activated cell sorting (FACS) tube, washed with 1x HBSS without Ca^2+^/Mg^2+^ (5 min, 1800 rpm, 21°C). Cells were then resuspended in complete RPMI cell media (RPMI, 10% heat inactivated fetal bovine serum, 50 μM 2-Mercaptoethanol, and 1% penicillin/streptomycin). Microglial cells were then counted using a hemocytometer and stained for flow cytometry analysis or exposed to lipopolysaccharide (LPS; see LPS immune challenge).

A subsample of isolated microglial cells (2×10^5^ cells/well) from each animal were transferred to individual wells of a 24-well plate containing complete RPMI and incubated overnight (12 hours at 37°C with 5% CO_2_) with LPS (40 ng/mL; *Escherichia coli* O127:B8 from Sigma-Aldrich, Burlington, MA, USA). Following this incubation period, wells were centrifuged (5 min, 400 x g) to pellet the cells, and the supernatant collected and stored at -80°C for later cytokine analysis.

### Splenocyte isolation

Whole spleens were homogenized in GentleMACs tubes (Miltenyi Biotec, Germany) while suspended in incomplete RPMI. The lysate was then transferred through a 70 μm cell strainer and washed with 2 mL of complete RPMI medium (RPMI, 10% heat inactivated fetal bovine serum, 50 μM 2-Mercaptoethanol, and 1% penicillin/streptomycin). Cells were pelleted (3 min, 800 x g) and supernatant was aspirated out. The cell pellet was disrupted, and 9 mL of sterile deionized water (4°C) was added to the cells for 8 seconds to lyse red blood cells, followed immediately by addition of 1 mL of 10x phosphate buffered saline (4°C). Cells were pelleted again (3 min; 800 x g) and supernatant aspirated before the cells were resuspended in 1 mL of complete RPMI cell medium. Splenocyte cells were then counted using a haemocytometer and Trypan Blue Stain (0.4%) to determine cell viability.

### Flow cytometry of microglia and splenocytes

Microglial cells were stained for flow cytometry by first washing in FACS buffer (phosphate buffered saline, 0.5% bovine serum albumin, and 2mM ethylenediaminetetraacetic Acid (EDTA), pH 7.2), followed by the addition of BV605 viability stain (0.1%) (BD Biosciences, Franklin Lakes, NJ, USA) for 15 min, 21°C. Cells were washed using FACS buffer and the supernatant decanted. Single stain controls were made using splenocytes for both antibody surface markers cluster of differentiation (CD)45-FITC and CD11b-APC Cy7 (BD Biosciences). Microglial samples were also stained with both CD45 and CD11b (BD Biosciences). Microglia and single-stain splenocyte controls were incubated in the dark for 30 minutes on ice. Cells were washed again, supernatant decanted, and the cells resuspended in paraformaldehyde (1%) and stored (4°C) until being sorted on the flow cytometer. Samples were run on a BD LSR II machine (BD Biosciences, USA) by the Flow Cytometry Core Facility, University of Calgary.

Splenocytes were stained for flow cytometry analysis with various surface and intracellular markers: PE-Cy7 anti-mouse CD19 (BD Biosciences), PerCP-Cy5.5 anti CD11b (BD Biosciences), APC-H7 anti-mouse CD8a (BD Biosciences), Alexa Fluor 700 anti-mouse CD3 (BD Biosciences), V500 anti-mouse CD3 (BD Biosciences), Brilliant Violet 421 anti-mouse CD25 (BD Biosciences), Brilliant Violet 421 anti-mouse CD11c (BD Biosciences), APC anti-mouse CD103 (BD Biosciences), Alexa Fluor 488 anti-mouse I-A/I-E (BD Biosciences), PE anti-mouse F4/80 (BD Biosciences), PE anti-gata3 (BD Biosciences), PE anti-mouse IL-17A (BD Biosciences), PE anti-mouse IL-12/23 (BD Biosciences), Alexa Fluor 488 anti-mouse IL-4 (BD Biosciences), Alexa Fluor 488 anti-mouse FoxP3 (BD Biosciences), Alexa Fluor 488 anti-mouse INF-γ (BD Biosciences), APC anti-mouse IL-6 (BD Biosciences), APC anti-mouse IL-10 (BD Biosciences), and eFluor 660 anti-mouse IL-13 (eBioscience, Thermo Fisher Scientific). Samples were incubated overnight in the dark (4°C) and stored in the dark (4°C) until being sorted on the flow cytometer. Samples were run on a CytoFLEX LX Flow Cytometer (Beckman Coulter, Brea, CA, USA) by the Flow Cytometry Core Facility at the University of Calgary.

Splenocyte flow cytometry samples were analyzed using FlowJo v10.6.2 software (FlowJo, Ashland, OR, USA) to gate for both cell type and cytokines based on intracellular and surface markers. Flow cytometry gating strategies were followed according to van Tilburg Bernardes et al.^28^ (**Supplementary Figure S1**). Splenocyte cell type profile doublets and dead cells were removed before gating based on the specific cell surface markers to determine cell phenotype (**Supplementary Figure S1**). Similarly, cytokine analysis was based on percentages of total live cells and doublet and dead cell elimination (**Supplementary Figures S2**).

### Cytokine measurements in colon, microglia, and lung tissue

Frozen colon and lung tissue samples were examined for cytokine and chemokine concentrations using the combination of two 96-well multiplex kits from Mesoscale Discovery (MSD; Rockville, MD. USA) including, V-PLEX Cytokine Panel 1 (mouse) (Cat# K15245D), and Proinflammatory Panel 1 (mouse) (Cat# K15048D) assays. Prior to assay determination, 50 − 150 mg of sample were homogenized in 1 mL of lysis buffer (150 mM NaCl, 20 mM Tris, 1 mM EGTA, 1% Triton X-100, protease inhibitor) for 4 min at 20 Hz using a tissue homogenizer (TissueLyser II, Qiagen). Homogenized samples were then centrifuged at 14,000 x g for 10 min to pellet debris, and supernatant collected and aliquoted to determine total protein corresponding concentrations by the Pierce BCA Protein Assay Kit (Thermo Fisher Scientific, Product No. 23225), with the rest used for MSD measurements. Acquired MSD data for each sample was then normalized to its total protein concentration prior to statistical analysis.

We also measured inflammatory markers in cell culture supernatants collected from the microglia LPS immune challenge, using the same MSD kits described above; however, we did not require protein determination per sample, as directed by the manufacturer.

### Ovalbumin (OVA)-induced allergic lung inflammation model

An experimental murine OVA model was followed as previously described^29^ with modifications (**Figure 1**). Mice were systemically sensitized intraperitoneally (IP) with sterile 200 μg of grade V OVA (Sigma-Aldrich) and 1.3 mg of aluminum hydroxide (Thermo Fisher Scientific) at weeks 7 and 9 post-birth. Following systemic sensitization, airway inflammation was induced by intranasal challenges at week 10 with 50 μg of Low Endo OVA (Whortington Biochemical, Lakewood, NJ, USA) for 3 consecutive days, followed by 2 days of 100 μg of grade V OVA (Sigma-Aldrich) in sterile phosphate-buffered saline (PBS). Intranasal challenges were done in mice under anesthesia with isoflurane. Following the challenge scheme, mice at P80 were anesthetized with ketamine (200 mg/kg; Vetoquinol, Lavaltrie, QC, Canada) and xylazine (10 mg/kg; Bayer Inc., Mississauga, ON, Canada), and BALF was collected by washing lungs with 3 x 1 mL of PBS + 10% fetal bovine serum (FBS) for total and differential cell counts. A hemocytometer was used to determine total BALF counts, while cell differentials (i.e., eosinophils, neutrophils, macrophages, and lymphocytes) were determined by counting 200 hematoxylin- and eosin-stained cells on a CytoSpin (Shandon Cytospin II, Thermo-Shandon, Runcorn, Cheshire, UK), and using standard cell morphological criteria to distinguish between cell types.

**Figure 1:**
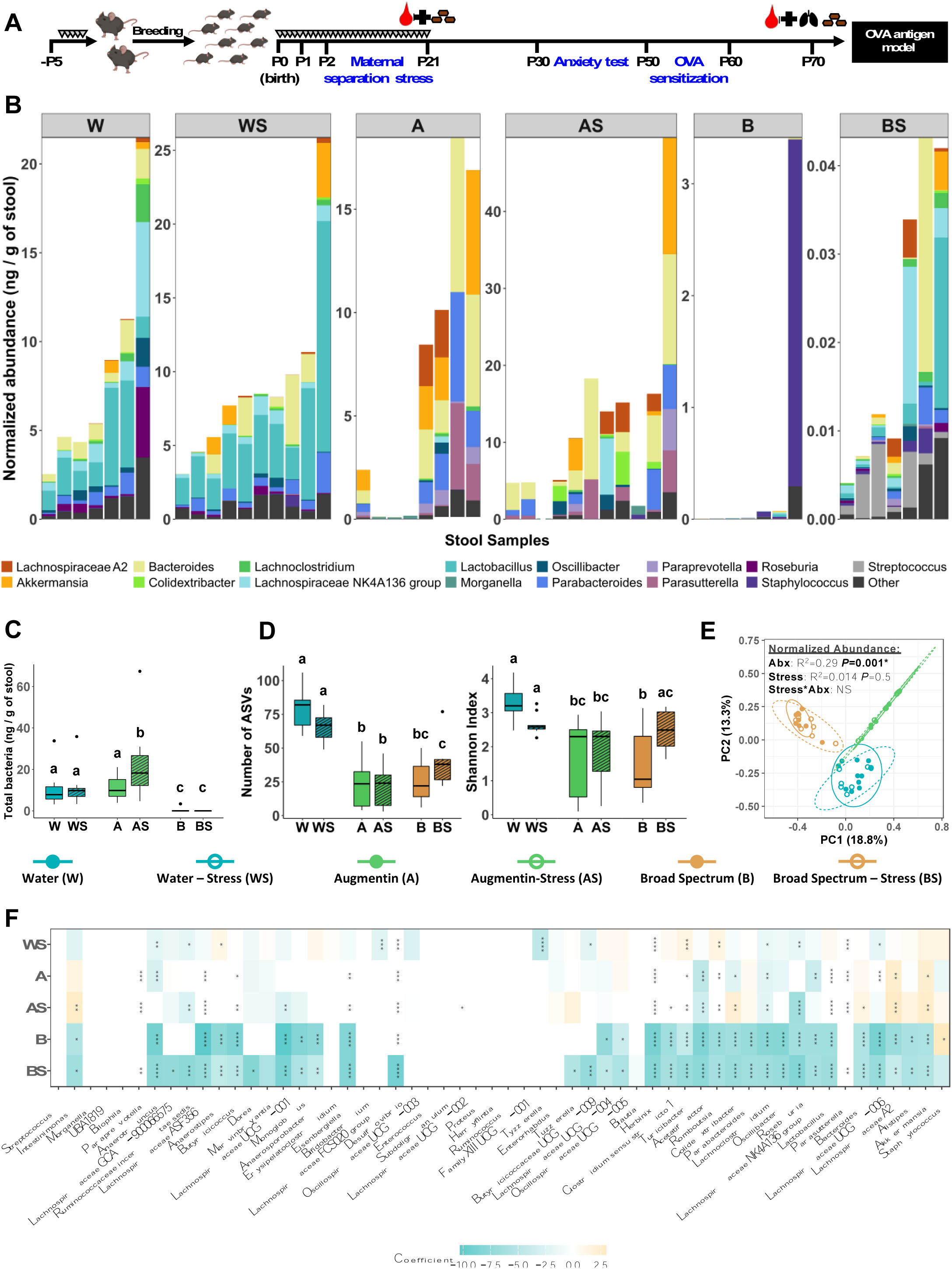
Early-life fecal microbiome characterization by stress and antibiotic treatments at P21. (A) Schematic diagram of the experimental design. C57BL/6 mice pups were exposed to antibiotics (broad spectrum antibiotic cocktail or Augmentin), or water as a control, by nursing on mothers provided with antibiotic treatments in drinking water, while simultaneously subjected to maternal separation stress protocols for up to 3 h/day (or left undisturbed as a control) from P0 to P21. Tissues were collected from pups at P21 with those remaining in the experiment behaviorally tested at P50 and later subjected to the OVA antigen lung inflammation model at P80. (B) Normalized abundance of the top 15 most predominant genera on a per sample basis across the treatment groups. (C) Total bacteria as determined by qPCR and (D) alpha-diversity (number of ASVs and Shannon index) across the treatment groups. Different lower-case letters denote significant differences between treatment groups (two-way ANOVA, p< 0.05, followed by a Fisher LSD test). (E) Beta-diversity using Bray-Curtis dissimilarity of normalized abundances at the ASV-level is visualized using principal coordinate analyses (PCA). Ellipses represent 95% confidence levels, and R^2^ and P values are from adonis models adjusted for antibiotic treatment, stress treatment, and an interaction term between antibiotics and stress; non-significant interactions were removed (denoted by NS) from the final models. (F) Compound Poisson linear regression models were used to determine differences in the normalized abundance of genera across treatment groups using MaAsLin2. Colours within each cell correspond to the model coefficient, with the water unstressed group as a reference. \**P*_FDR_<0.10, \*\**P*_FDR_<0.05, \*\*\**P*_FDR_<0.01, \*\*\*\**P*_FDR_<0.001. Water (W); Water-Stress (WS); Augmentin (A); Augmentin-Stress (AS); Broad spectrum antibiotic (B); Broad spectrum antibiotic-Stress (BS).

### Behavioural assessment

The elevated plus maze (EPM) was used to assess animal behaviour at P60. All behavioural tests were performed in the morning, from 7 am to 12 pm. On the experimental day, mice were brought to the behavioural room and allowed to rest and habituate for at least 2 h prior the test. All analyses were done by an experimenter blinded to the treatment groups. For the EPM test, mice were randomly selected and placed on the EPM apparatus. The apparatus consisted of a cross-shaped platform with an acrylic surface elevated 50 cm above the floor. The 4 arms were composed of two opposite open arms with no walls and two enclosed arms by a 20 cm wall, all connected to a central area of 7 × 7 cm. Each arm was 21.5 cm long and 7 cm wide. Light intensity was set to 70−75 lux on the open arms and 15−20 lux on the closed arms. The recording camera was set above the apparatus. In each session, the animal was placed on the centre zone facing the open arm opposite to the experimenter and recorded for 5 min. At the end of the session, the subject mouse was returned to a new home cage. The apparatus was cleaned with 70% ethanol in between subject mouse. TopScan software (CleverSys, Reston, VA, USA) was used for analyses of the recorded videos, assessing the animals time spent in the open and closed arms, and total locomotion through the maze.

## Bioinformatic and data analyses

### Pre-processing, analysis, and visualization of microbiome data

Bioinformatics and data analyses were conducted in R (version 4.2.3, R Core Team, 2017). Sequencing data was quality-checked, trimmed, merged, and chimeras were removed using DADA2 version 1.24.0.^30^ Amplicon sequence variants (ASVs) were assigned taxonomy using the SILVA version 138 database.^31^ Raw sequences have been deposited in the NCBI database under BioProject accession number PRJNA1204146.

ASVs identified in negative DNA extraction and PCR controls were removed from the P21 and P80 datasets separately (n=15 and n=4 ASVs, respectively). Phyloseq (version 1.44.0) was then used for downstream data processing and analysis.^32^ Singleton and doubleton ASVs were removed across the entire dataset, in addition to ASVs belonging to eukaryotes, chloroplasts, and mitochondria. Of note, other ASVs mapping to Cyanobacteria were retained as they aligned to Vampirivibrionia which is non-photosynthetic. Data was rarefied (1,420 reads for P21; 28,981 reads for P80) according to rarefaction curves and minimum sequence coverage, and samples with read counts below the cut-off were excluded (n=2 samples for both datasets). At the early-life timepoint (P21), a total of 50 samples were included with a total of 367 unique ASVs; for the later timepoint (P80), a total of 56 samples and 857 ASVs were included in the analyses.

Normalized abundances of ASVs were also calculated by multiplying the ASV relative abundances by the total bacteria (ng) per gram of stool (determined from qPCR) for each sample, which resulted in the normalized abundance of each ASV expressed in the units, ng of bacteria per gram of stool. Phyloseq was then used to calculate and plot microbial relative and normalized abundances (phylum and genus levels), alpha-diversity (number of ASVs, Shannon index), and beta-diversity (Bray-Curtis dissimilarity). The ggplot2 (version 3.5.0)^33^ and vegan (version 2.6-4)^34^ packages were also used for data visualizations.

### Statistical analyses

GraphPad Prism v.8.4.1 was used to compare the following continuous outcomes between treatment groups: microbial measures (total bacteria, number of ASVs, Shannon Index), percent activated microglia, plasma and hair (body hair and whiskers) corticosterone concentrations, cytokine concentrations (gut, lung, and microglia, splenocytes), SCFA concentrations, lung inflammatory cell counts, and elevated plus maze behavioral data. In each case, a two-way ANOVA was used to assess differences between groups including a Fisher least significance difference (LSD) test for treatment group comparisons (p<0.05). Data was visualized as boxplots using the ggplot2 R package.

Differences in the prevalence (i.e., presence/absence) of microbial genera between treatment groups was assessed using Fisher’s Exact Tests; differences in the relative and normalized abundance of microbial genera between treatment groups was determined using compound Poisson linear regression models in MaAsLin2 version 1.14.1.^35^ These analyses were conducted separately for the P21 and P80 time points, and were visualized in heatmaps generated using ggplot2.

To determine whether antibiotic treatments could be predicted from the P21 microbial data, random forest models were built with 30 trees (selected to minimize error rates) using normalized genera abundances.^36^ Of note, preliminary random forest models were also constructed using all treatment groups (antibiotic and stress combinations); however, the models were unable to distinguish between the stressed and unstressed groups and, therefore, were combined within the antibiotic treatments to improve accuracy of the final model. ASVs with the greatest mean decrease in Gini values were visualized to indicate their weighted importance in predicting the antibiotic groups.

To integrate the multi**-**system and multi**-**omic nature of our data, we conducted a multi-block partial least squares-discriminant analysis (PLS-DA), referred to as DIABLO, using mixOmics (version 6.24.0).^37^ Models were constructed using P21 data including microglia (cytokines, activation profile), normalized microbial genera abundance, corticosterone (plasma, body hair and whiskers), colon (cytokines), splenocytes (cytokines and cell populations), and SCFAs. Correlation circle and Circos plots were then used to visualize the model results. Of note, similar to the random forest analyses, only antibiotic treatments were tested rather than all treatment groups (antibiotic and stress combination) to improve model accuracy.

## Results

### Antibiotic and stress-induced disruptions of the early-life and adult gut microbiome

Exposing newborn mice to the six treatment combinations of antibiotics (water, Augmentin, broad spectrum) with or without stress resulted in significantly distinct microbiome shifts (**Figure 1**). To account for the expected changes in bacterial load from antibiotic treatments, we normalized the abundance of each ASV to the total bacterial DNA amount per g of stool on a per sample basis. Relative and normalized abundances of the top 15 genera per sample can be found in **Supplementary Figure S3A and Figure 1B**, respectively. Importantly, given the range of bacterial loads observed in each sample, normalized abundances allowed for more accurate comparisons of bacterial amounts between samples − otherwise findings would be masked when considering only relative abundances. When comparing treatment groups, total bacterial concentrations in Augmentin-treated animals increased (when in combination with stress), while those treated with the broad spectrum antibiotic cocktail decreased, compared to water-unstressed animals (**Figure 1C**). Significant microbiome changes were also reflected with a decrease in alpha-diversity (number of ASVs and Shannon index) at P21 in both antibiotic-treated groups, particularly in the broad-spectrum antibiotic treated mice where the bacterial community was nearly eliminated (**Figure 1C-D**). These antibiotic-dependent effects resulted in significant changes in the overall microbial community structure using Bray-Curtis dissimilarity of normalized abundances (**Figure 1E**). Most notably, significant shifts in the first two principal components (PC1 and PC2) were largely explained by antibiotic treatment (R^2^ = 0.29; *P* = 0.001) with distinct clusters generated for each group, while the effects of early-life chronic stress did not significantly contribute to changes in microbial beta-diversity.

The prevalence (presence/absence), relative, and normalized abundances of individual genera changed considerably across the different treatment groups (**Figure 1F**, **Supplementary Figure S3C**). More specifically, the low concentrations of bacteria in the broad spectrum-treated mice resulted in many differentially-abundant genera, demonstrating significant decreases in key commensal genera involved in carbohydrate fermentation such as, *Lactobacillus* (*P* False Discovery Rate _(FDR)_<0.001), *Bacteroides* (*P*_FDR_<0.001), *Lachnospiraceae* (NK4A136 group) (*P*_FDR_<0.001), *Parabacteroides* (*P*_FDR_<0.001), and *Roseburia* (*P*_FDR_<0.001; **Figure 1F**). Similar decreases in genera were observed in Augmentin-treated mice, although shifts were less prominent (*P*_FDR_<0.05−0.01) than in the broad spectrum-treated mice. Interestingly, *Intestinimonas* (*P*_FDR_<0.05), *Staphylococcus* (*P*_FDR_<0.1), *Streptococcus* (*P*_FDR_<0.05), *Bacteroides* (*P*_FDR_<0.1), and *Akkermansia* (*P*_FDR_<0.1−0.05) concentrations were significantly increased in Augmentin-treated mice compared to the water-unstressed group, suggesting that this antibiotic treatment favoured the overgrowth of these anaerobic bacterial species, which are conventionally targeted by Augmentin. Finally, we used a Random Forest classifier to determine if microbiome shifts (normalized abundances of individual ASVs) could be predicted from the antibiotic treatments. The model strongly classified antibiotic grouping with a 98% accuracy, with only 1 misclassification of an Augmentin sample predicted to be from the broad spectrum group. An ASV mapping to *Parabacteroides* was the most predictive feature in this model, followed by ASVs mapping to *Romboutsia, Lactobacillus,* and Lachnospiraecae NK4A136 group (**Supplementary Figure S3**).

Compared to the observations in antibiotic-treated mice, the effects of maternal separation-induced stress on the microbiome were very reduced. Nevertheless, at P21, stress increased total bacterial concentration in mice treated with Augmentin compared to untreated mice, and it mitigated the loss of alpha-diversity in mice treated with broad-spectrum antibiotics (**Figure 1C-D**).

To determine the long-term effects of early-life antibiotic treatments, we analyzed the microbiome at P80. Intriguingly, differences in relative and normalized abundances, total bacteria concentration, and alpha- and beta-diversity persisted and, in some cases, were even more apparent in adulthood, demonstrating the lasting effects of early-life antibiotics on the microbiome (**Supplementary Figure S4**). Furthermore, the presence (prevalence) of key commensal taxa at P80, including *Lactobacillus*, *Mucispirillum*, *Lachnospiraceae*, *Roseburia*, *Intestinimonas*, *Oscillibacter*, *Acetatifactor*, and *Tyzzerella* were drastically and significantly reduced in broad spectrum-treated mice, and to a lesser extent in Augmentin-treated mice, and had a greater number of significantly reduced taxa than was observed at P21, compared to the water-unstressed group (**Supplementary Figure S5**). In contrast, the presence of other common gut microbes such as *Enterococcus* and *Blautia* increased in antibiotic groups. Numerous changes in bacterial relative and normalized abundances were also observed at P80, with similar taxa showing differential abundances across the treatment groups as those observed with the prevalence analyses (e.g., increased abundances of *Blautia* across antibiotic and/or stress groups, compared to water-unstressed; **Supplementary Figure S4 and S5**). Of note, early-life stress alone did not result in persistent microbiota differences at P80 (**Supplementary Figure S4 and S5**).

Taken together, these results highlight that early-life antibiotic exposure has profound and lasting impacts on the gut microbiome, with broad-spectrum antibiotics inducing more severe disruptions than Augmentin. These findings underscore the importance of normalization approaches for accurate microbiome analysis and reveal a minimal role for maternal separation-induced stress in shaping microbial composition during early life or adulthood.

### Short-chain fatty acid stool concentrations in early-life and adulthood

Given the observed decrease in fermentative taxa in antibiotic-treated mice, we determined the concentration of SCFAs as main by-products of microbial carbohydrate fermentation in the colon. Consistent with the microbiome analysis findings, SCFA concentrations were drastically decreased at 21-days postnatally in both antibiotic-treated groups, but more prominently in broad spectrum-treated mice (**Figure 2A**). Butyrate, isobutyrate, isovalerate, valerate, propionate, and acetate were all below detection limits in the broad-spectrum group. While some of these SCFAs were detectable in Augmentin-treated mice, they were still significantly decreased compared to both water control groups (stressed and unstressed) (**Figure 2A**). Intriguingly, early-life stress induced a significant decrease in butyrate in mice not treated with antibiotics, indicating that while stress has a lesser effect on the microbiota at the taxonomic level, it induced functional changes to microbial fermentation by-products.

**Figure 2:**
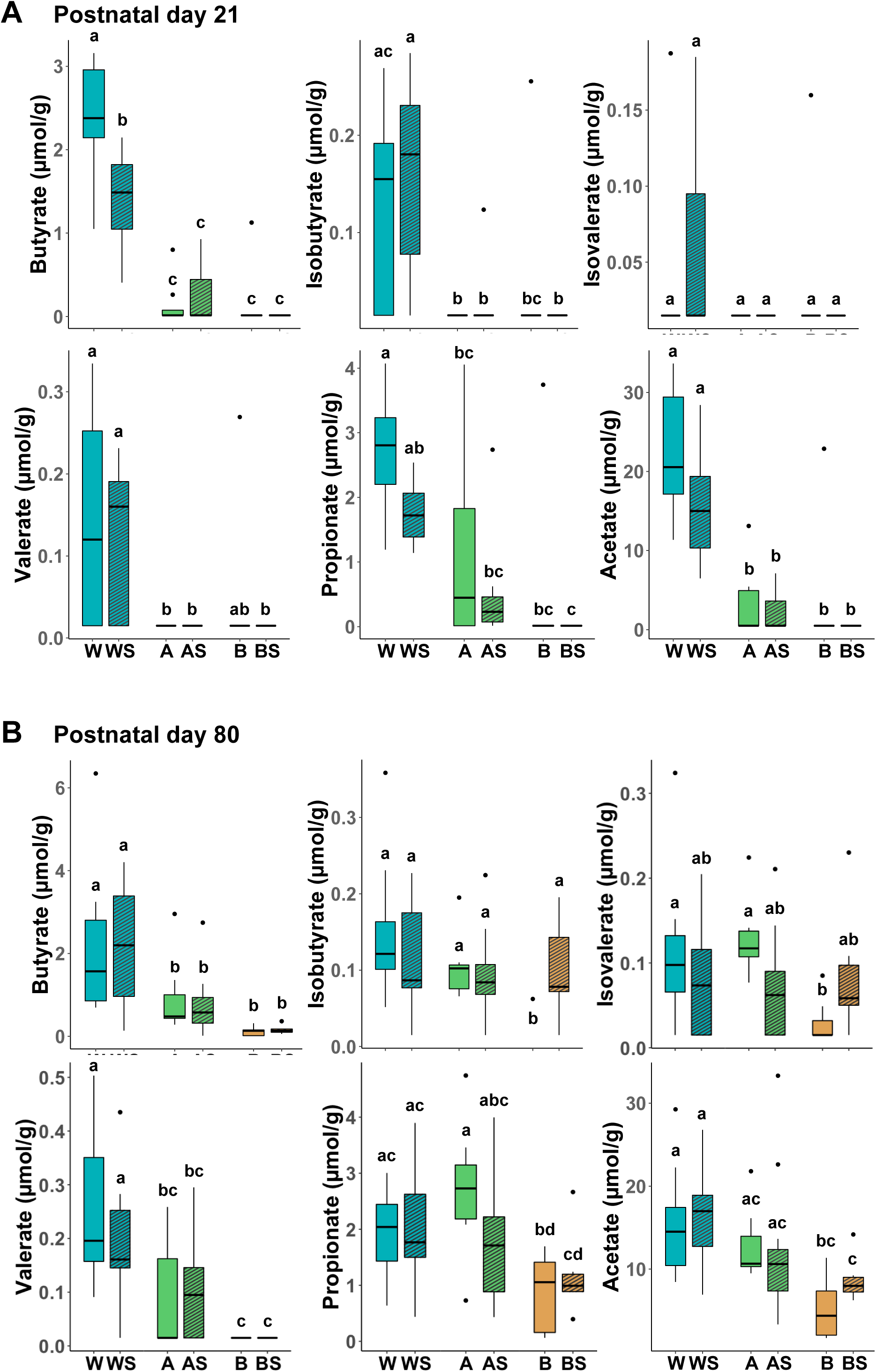
Stress and antibiotic-induced modulation of fecal short-chain fatty acid concentrations in early-life and adulthood. Short chain fatty acid (SCFA) analysis in stool from mouse pups at (A) postnatal (P) day 21 and (B) P80. Different lower-case letters denote significant differences between treatment groups (two-way ANOVA, p<0.05, followed by a Fisher LSD test). Water (W); Water-Stress (WS); Augmentin (A); Augmentin-Stress (AS); Broad spectrum antibiotic (B); Broad spectrum antibiotic-Stress (BS).

While the levels of SCFAs at P80 had recovered and were detectable in all treatment groups, they remained significantly lower in broad spectrum-treated mice (stressed and unstressed) for butyrate, valerate, propionate, and acetate, and in unstressed mice for isovalerate and isobutyrate when compared to the water-treated unstressed mice (**Figure 2B**). Similarly, SCFA concentrations for butyrate and valerate were also lower in Augmentin-treated mice (stressed and unstressed), although the concentrations of propionate and isovalerate were highest in the unstressed mice compared to the other treatment conditions.

Overall, these results show that early-life antibiotic treatment induces a vigorous acute depletion of fermentative metabolism, which can persist into adulthood when the early-life microbiome is perturbed with broad spectrum antibiotics.

### Microglial phenotype and immunogenicity in early-life

Activated microglia profiles were analyzed using flow cytometry by comparing the proportion of CD11b+CD45mid live cells at P21 across groups (see staining strategy in **Supplementary Figure S6**). Compared to untreated mice, the proportions of activated microglia were significantly higher in mice treated with combinations of Augmentin and stress, and broad spectrum antibiotics and stress (**Figure 3A**). Augmentin-stress exposed mice also had significantly elevated proportions of activated microglia in comparison to mice exposed solely to broad spectrum antibiotics, or stress alone. In general, stress exposure resulted in observable but non-significant trends of increased microglial activation within each antibiotic treatment group.

**Figure 3:**
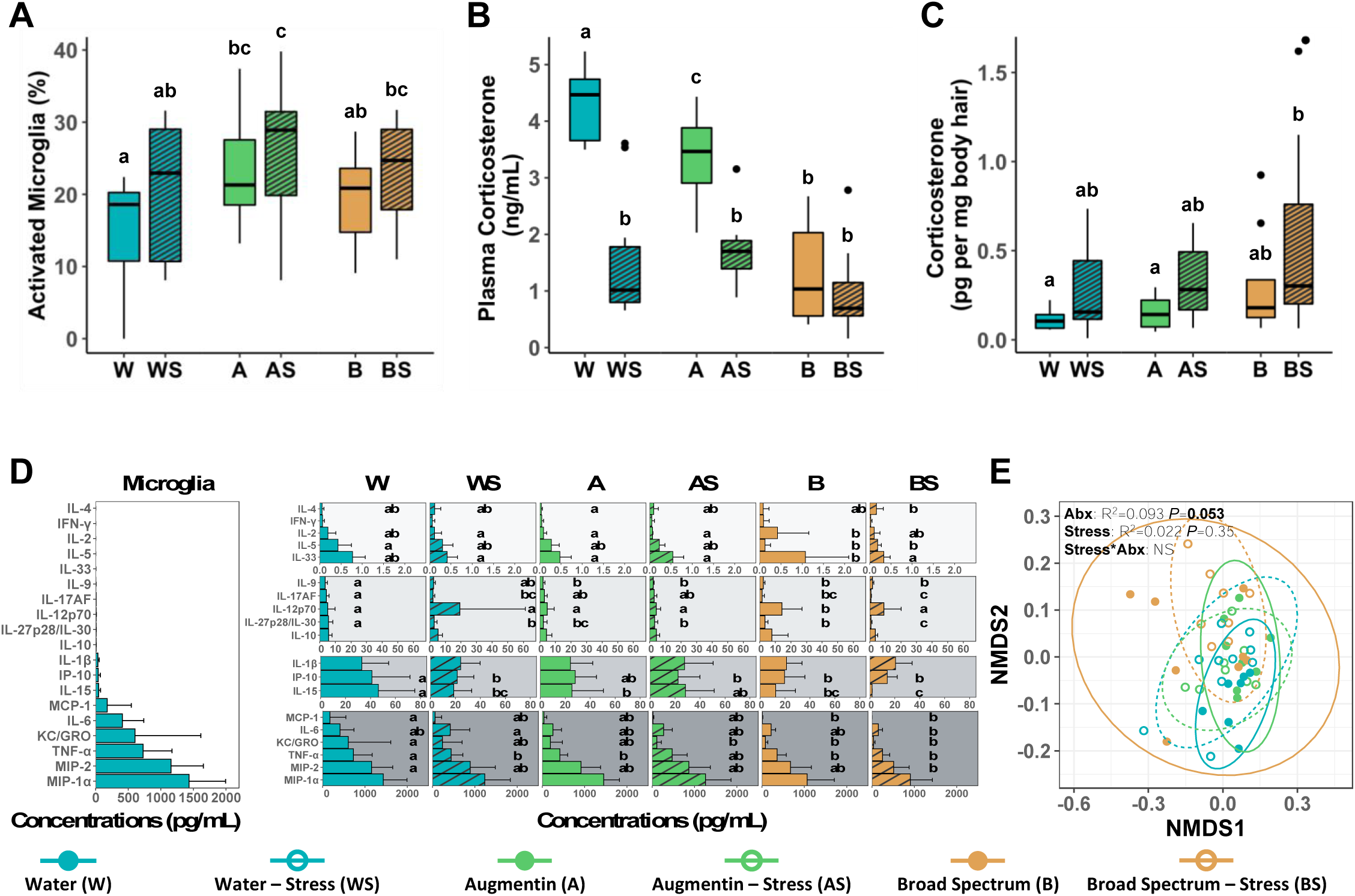
Early-life responses to stress and antibiotic treatments in microglial activation and corticosterone metabolism at P21. (A) Tissues were assessed for the percentage of activated brain microglia using fluorescence-activated cell sorting analysis. Corticosterone levels were assessed in (B) plasma and (C) body hair using ELISAs. (D) An in vitro analysis of secreted cytokines from LPS-stimulated microglia cells was conducted using the Mesoscale Device (MSD) electrochemiluminescence multiplex assays. Microglia cytokine concentrations are displayed across a single concentration range on the left panel to show large variation in marker levels in the control group (water-unstressed), while organized on the right panel into discrete cytokine concentration ranges for visual clarity. Different lower-case letters denote significant differences in cytokine concentrations between treatment groups; letters are only displayed for cytokines with statistically significant findings (two-way ANOVA, p < 0.05, followed by a Fisher LSD test). (E) Non-metric multidimensional scaling (NMDS) ordination plot of cytokine concentrations is shown. Ellipses represent 95% confidence intervals, and R^2^ and P values are from adonis models adjusted for antibiotic treatment, stress treatment, and an interaction term between antibiotics and stress; non-significant interactions were removed (denoted by NS) from the final models. Water (W); Water-Stress (WS); Augmentin (A); Augmentin-Stress (AS); Broad spectrum antibiotic (B); Broad spectrum antibiotic-Stress (BS).

To further study this, isolated microglia were exposed overnight to bacterial LPS. Microglia from mice treated with broad spectrum antibiotics alone and/or in combination with stress, had reduced cytokine production compared to the water unstressed group (**Figure 3D**). This included significant decreases in interleukin (IL)-5, IL-6, IL-9, IL-17AF, IL-27p28/IL-30, IL-15, as well as IP-10 and tumor necrosis factor (TNF)-a. Interestingly, many chemotactic cytokines related to cell activation were significantly decreased in not only the broad-spectrum group, but also in Augmentin-treated and water-fed mice exposed to chronic stress. These included monocyte chemoattractant protein-1 (MCP-1), keratinocyte chemoattractant/human growth regulated oncogene (KC/GRO), and macrophage inflammatory protein-2 (MIP-2), which were among the highest secreted cytokines detected in LPS-exposed cultured microglia (**Figure 3D**). An ordination analysis of cytokine concentrations using non-metric multidimensional scaling (NMDS) showed that exposure to antibiotics shifted the overall cytokine production of microglia upon exposure to LPS, although the result was marginally significant (*P* = 0.053; **Figure 3E**). Similar to the microbiome results, our findings showed that early-life antibiotic treatment, and to a lesser extent, stress, induced potent reductions in cytokine and chemokine secretion in microglia.

### HPA stress axis activity and behaviour in early-life and adulthood

With respect to HPA axis activation, both antibiotic and stress exposure significantly decreased the basal circulating levels of plasma corticosterone in mice at P21, compared to the water unstressed group (**Figure 3B**). Stress in particular had strong additive effects on depressed corticosterone levels for both Augmentin and broad spectrum antibiotic treatments. Even in the absence of antibiotics, stress alone (with water) significantly lowered basal levels of corticosterone. In contrast, corticosterone accumulated at higher levels in body hair and whisker follicles of P21 mice exposed to both stress and antibiotic treatments (especially the broad-spectrum stressed group), compared to the water unstressed group (**Figure 3C, Supplementary Figure S7D**). Interestingly, these increased levels of body hair-accumulated corticosterone persisted in adult P80 mice that were chronically stressed as neonates, regardless of antibiotic treatment (**Supplementary Figure S7E**). Moreover, body hair collected from dams of the chronically-stressed pups did not show significant differences in corticosterone accumulation between treatment groups, despite a small increase in the broad spectrum-exposed mice (**Supplementary Figure S7F**), indicating that pup stress levels were likely independent of their mother’s.

In addition to measuring HPA stress axis activity in circulation and hair, we also tested mice at 8 weeks of age (P60) for behavioural changes and indications of anxiety traits that could result from the daily maternal separation stress prior to P21. Stress in combination with Augmentin explained an increased number of entries and percentage of time in the open arm test, suggesting increased exploratory behaviour in these mice; however, no other significant differences in open arm entries, time in the open arm, nor locomotion were observed between treatment groups (**Supplementary Figure S7**).

### Intestinal and systemic immune response in early-life

To determine the effect of early-life antibiotic treatment and stress on local intestinal and systemic immunity, we determined cytokine responses in colonic tissue using chemiluminescence detection (Mesoscale) and surveyed immune cell populations and cytokine production in splenocytes using flow cytometry. We detected a variety of changes in colonic tissue cytokine levels, including significant increases in pleotropic IL-9, INF-γ, IL-6, IP-10, IL-6, and IL-12p70 in the broad-spectrum stressed group, compared to the water-unstressed group. Similarly, IL-4, MIP-2, IL-1β were significantly increased in the Augmentin-unstressed group compared to the control group (**Figure 4A**). Ordination of overall cytokine responses did not show significant differences between tissue cytokines in any of the treatment combinations (**Figure 4B**).

**Figure 4:**
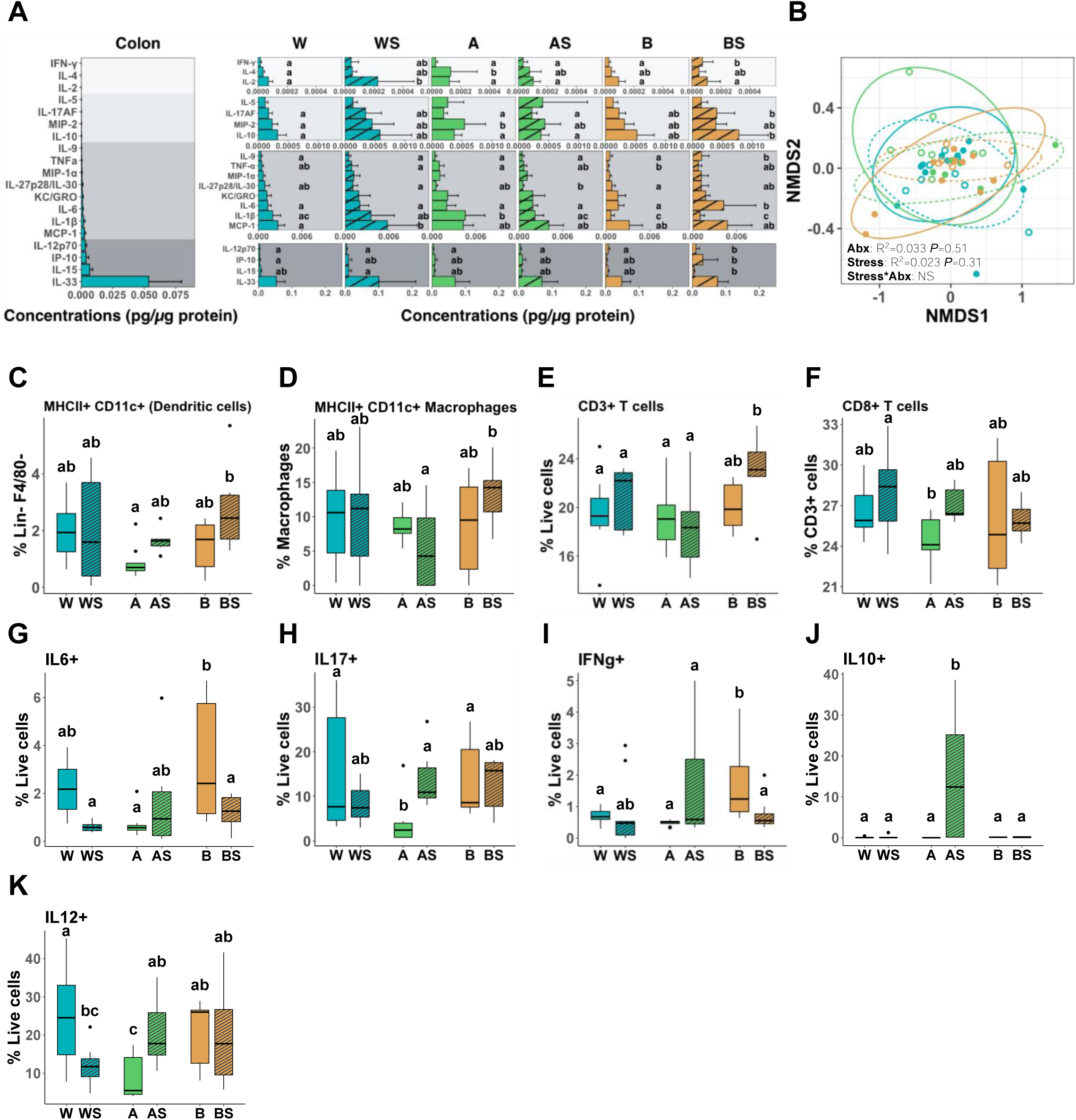
Early-life responses to stress and antibiotic treatments in systemic and intestinal immunity at P21. (A) Colon tissues from mouse pups were assessed for cytokine concentrations using the Mesoscale Device (MSD) electrochemiluminescence multiplex assays. Colon cytokine concentrations are displayed across a single concentration range on the left panel to show large variation in marker levels in the control group (water-unstressed), while organized on the right panel into discrete cytokine concentration ranges for visual clarity. (B) Non-metric multidimensional scaling (NMDS) ordination plot of colon cytokine concentrations is shown.

Flow cytometric analysis of isolated splenocytes identified a decrease in conventional dendritic cells (Lin^−^F4/80^−^ MHCII^+^ CD11c^+^) in the Augmentin unstressed group compared to broad spectrum stressed-exposed mice (**Figure 4C**), while Lin^−^F4/80^+^ MHCII^+^ CD11c^+^ macrophages were increased in the broad spectrum stressed group compared to the Augmentin stressed group (**Figure 4D**). Meanwhile, CD3^+^ T lymphocytes (T cells) were increased in broad spectrum-treated stressed mice (**Figure 4E**), and CD8^+^ T cells were decreased in Augmentin unstressed group, compared to water treated groups (**Figure 4F**). Other cell types measured in each treatment group can be found in Supplemental Figure S8.

In contrast to the discrete differences in immune cell populations, we detected larger differences in cytokine levels measured by flow cytometry (**Figure 4G-K; Supplemental Figure S8**). Broad spectrum antibiotics (without stress) significantly increased levels of IL-6, IL-17 and IFN-γ in all live cells compared to the Augmentin unstressed mice, while Augmentin and stress resulted in an increase of IL-10 compared to all other groups, and Augmentin alone decreased IL-12 compared to all other groups (**Figure 4G-K**). Overall, while the effects of antibiotic and stress could be detected in several responses in the colon and the spleen, these were less prominent than those observed in stimulated microglia.

Ellipses represent 95% confidence intervals, and R^2^ and P values are from adonis models adjusted for antibiotic treatment, stress treatment, and an interaction term between antibiotics and stress; non-significant interactions were removed (denoted by NS) from the final models. (C-K) Percentage of spleen cell populations and cytokine-producing splenocytes using FACS analysis. For panels A and C-K, different lower-case letters denote significant differences between treatment groups; letters are only displayed for cytokines with statistically significant findings (two-way ANOVA, p < 0.05, followed by a Fisher LSD test). Water (W); Water-Stress (WS); Augmentin (A); Augmentin-Stress (AS); Broad spectrum antibiotic (B); Broad spectrum antibiotic-Stress (BS).

### Allergic airway inflammatory response in adult mice

Seven week old adult mice were sensitized systemically and later challenged intranasally with OVA to induce an allergic immune response in the airways and determine the effect of early-life antibiotics and stress in lung immunity (**Figure 5A**). As expected, this model resulted in significantly elevated levels of inflammatory cells in all treatment groups compared to naive mice that were treated with a vehicle control (**Figure 5B**). Augmentin-treated mice in combination with stress, displayed the highest infiltrate of total inflammatory cells in the BALF, compared to mice given water or broad-spectrum antibiotics. Otherwise, stress did not influence allergic airway inflammation to OVA in other treatment combinations as there were no significant differences.

**Figure 5.**
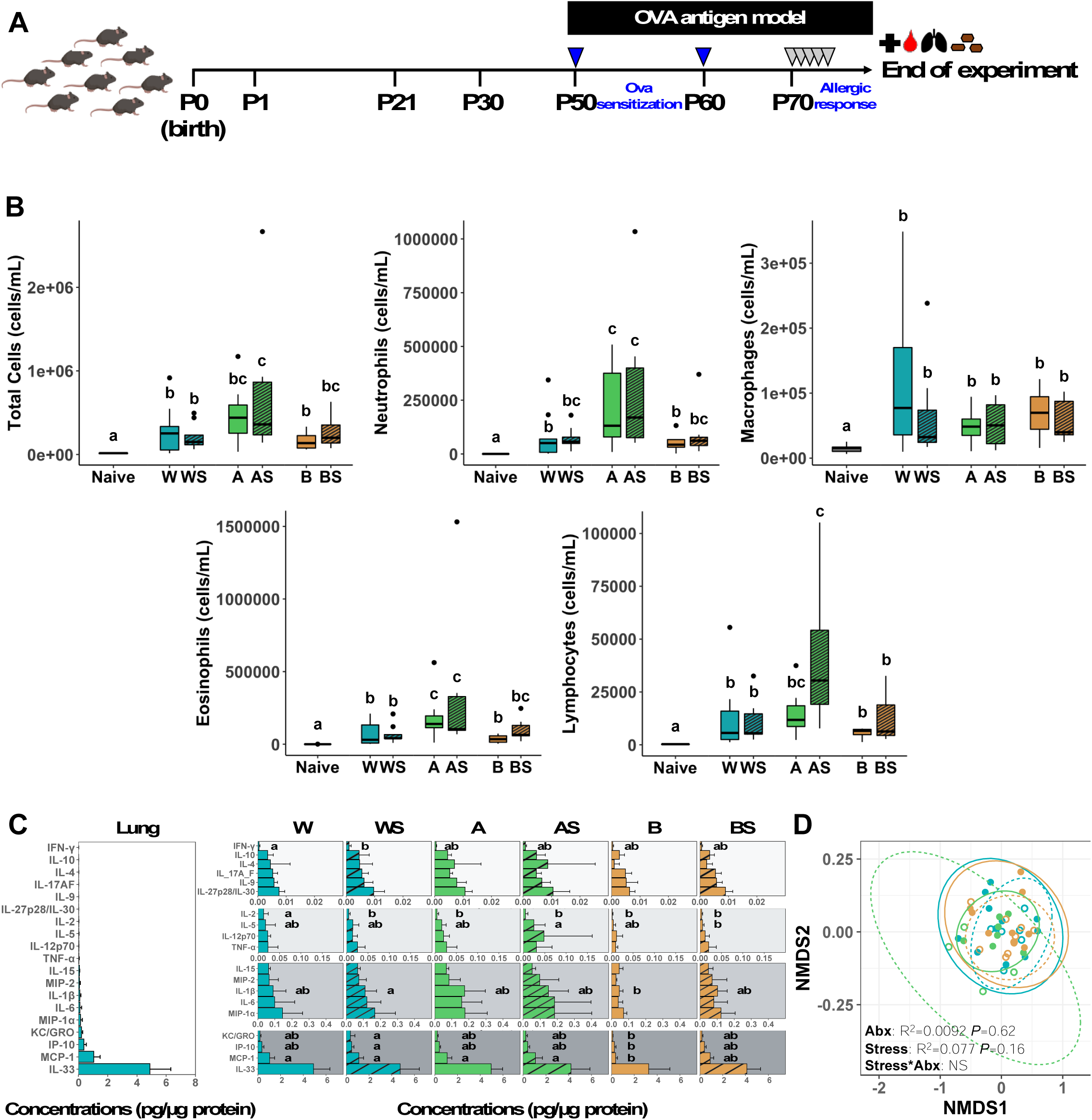
Allergic airway inflammatory responses to stress and antibiotic treatments at P80. (A) Schematic diagram of the OVA antigen model of allergic airway inflammation. Antibiotic and/or stress-exposed C57BL/6 mice were systemically sensitized with two intraperitoneal (*i.p.*) injections of OVA+Alum at weeks 7 and 9, and intranasally challenged with OVA for 5 consecutive days to stimulate lung inflammation. A group of mice were kept unchallenged as naive mice (see Methods). (B) Total cellular counts in bronchoalveolar lavage fluid (BALF) following OVA challenge with differential cell counts for neutrophils, macrophages, eosinophils, and lymphocytes in. (C) Cytokine concentrations measured in lung tissue lysates following the OVA antigen challenge using the Mesoscale Device (MSD) electrochemiluminescence multiplex assays. Cytokine concentrations are displayed across a single concentration range on the left panel to show large variation in marker levels in the control group (water-unstressed), while organized on the right panel into discrete cytokine concentration ranges for visual clarity. (D) Non-metric multidimensional scaling (NMDS) ordination plots of lung cytokine concentrations is shown. Ellipses represent 95% confidence intervals, and R^2^ and P values are from adonis models adjusted for antibiotic treatment, stress treatment, and an interaction term between antibiotics and stress; non-significant interactions were removed (denoted by NS) from the final models. For panels B-C, different lower-case letters denote significant differences between treatment groups within B-C (Total BALF: one-way ANOVA, p<0.05, followed by a Fisher LSD test; Lung MSD: two-way ANOVA, p<0.05, followed by a Fisher LSD test). Water (W); Water-Stress (WS); Augmentin (A); Augmentin-Stress (AS); Broad spectrum antibiotic (B); Broad spectrum antibiotic-Stress (BS).

Differential analysis of the principal immune cell types present in BALF by microscopy showed significantly elevated levels of neutrophils, eosinophils and lymphocytes in mice exposed to a combination of Augmentin and stress, when compared to the water-exposed mice. Lymphocytes in particular were significantly elevated compared to all other groups (**Figure 5B**).

Lung tissue cytokine levels were measured with limited significant differences observed in the targeted cytokines. IL-2 was the most significantly affected cytokine with decreased levels in all treatment groups compared to water unstressed mice, while IL-5 was also significantly decreased in the broad spectrum stressed group compared to the Augmentin stressed groups (**Figure 5C**). Of interest, while KC/GRO, IP-10, MCP-1 were all significantly decreased in the broad spectrum unstressed group compared to the water stressed group, MCP-1 was also significantly lower to all, but the broad spectrum stressed group. Finally, the overall ordination of cytokine profiles did not show significant differences in antibiotic or stress treatments (**Figure 5D**).

Altogether, only early-life Augmentin treatment induced changes in immune cell infiltrate levels that led to increased susceptibility to allergic lung inflammation in adulthood, while broad spectrum antibiotics significantly decreased inflammatory cytokine concentrations more than any other treatment group.

### Systems-level integration of microbial, immune, and HPA axis responses

To explore the complex interconnections among the biological processes assessed in our study, we conducted an integrative analysis using PLS-DA of microbial, SCFA, stress, and immune datasets at P21 (Figures 6 and 7). This method efficiently handles multi-omic datasets making it well-suited for identifying relationships between microbes, microbial by-products, and host biomolecules.^37,38^

**Figure 6:**
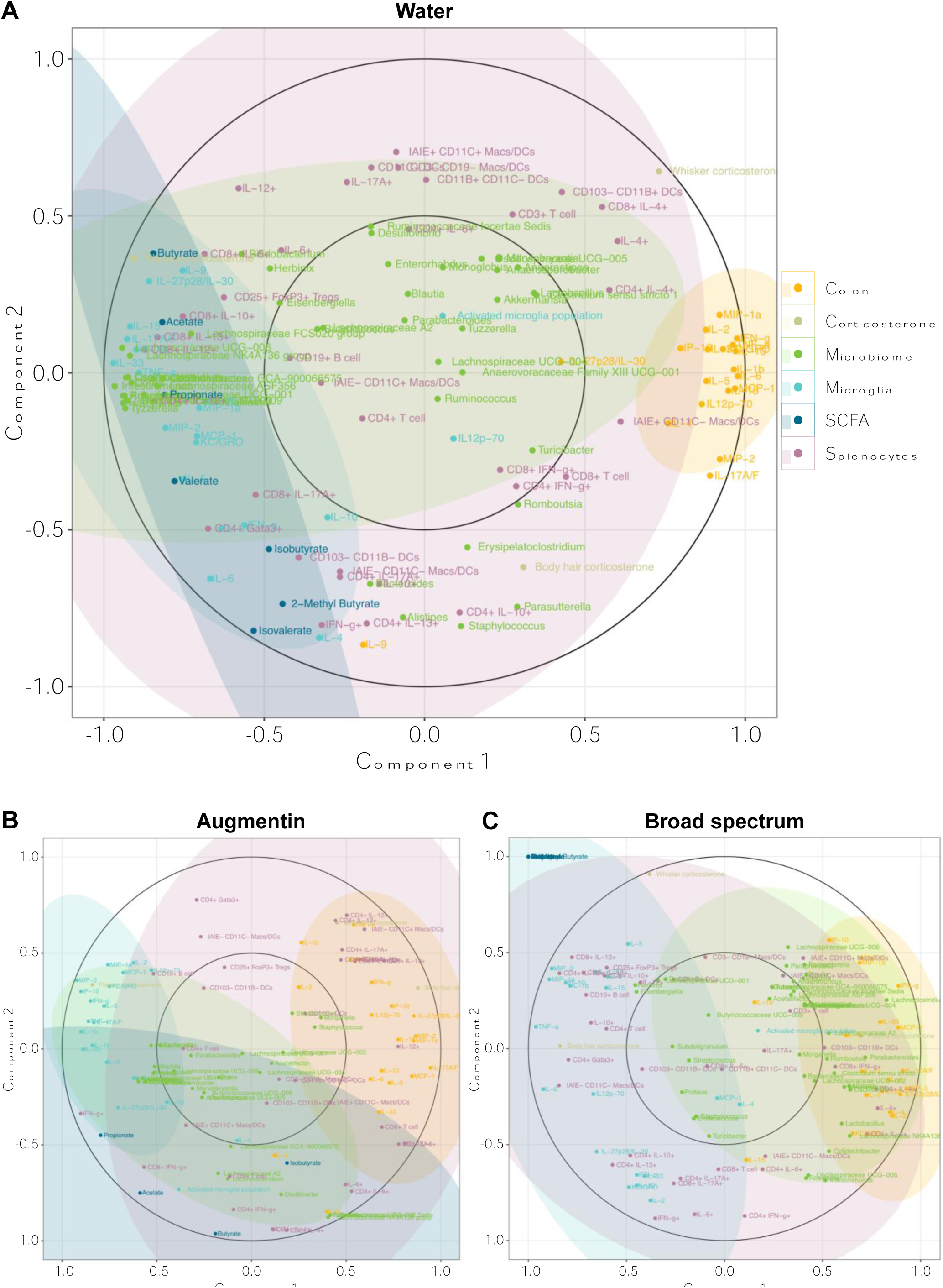
Systems-level integration of microbial, immune, and HPA axis responses using a multi-block PLS-DA analysis in postnatal day 21 mice treated with antibiotic and stress treatments. A correlation circle plot was used to visualize the different variable types (i.e., colon, corticosterone, microbiome, microglia, SCFAs, and splenocytes) across the first and second principal components for (A) water, (B) Augmenting and (C) broad spectrum antibiotic treatments. Colours correspond to the different variable types, and individual data points between the inner and outer circles (between ±0.5 and ±1) represent stronger associations along each component.

**Figure 7.**
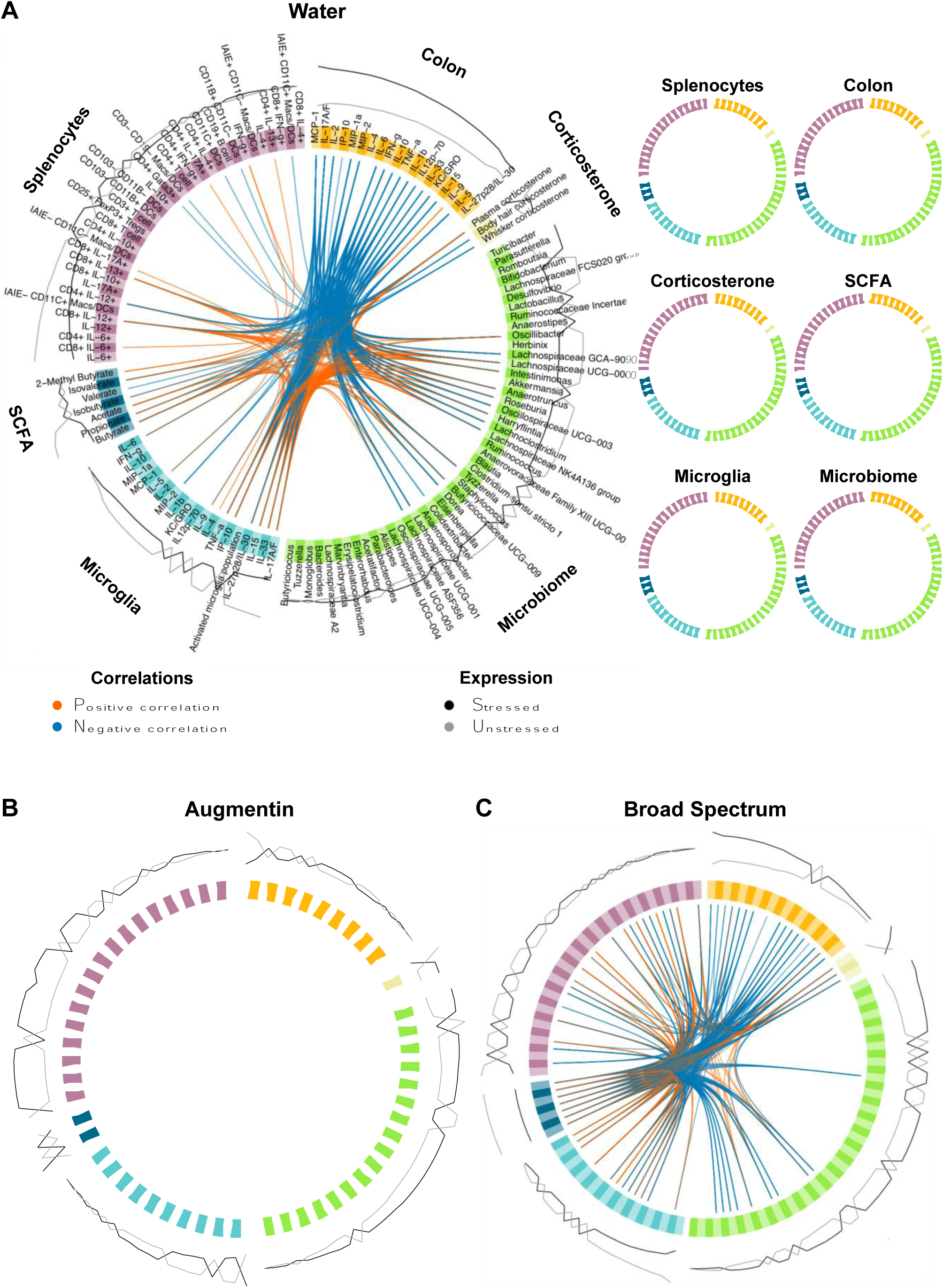
Systems-level integration of microbial, immune, and HPA axis responses at P21 in mice exposed to antibiotic and stress treatments. Circos plots generated for (A) water, (B) Augmentin and (C) broad spectrum antibiotic treatments from a multiblock PLS-DA analysis in mixOmics show correlations between individual microbial, immune, and HPA axis variables. Positive (red) and negative (blue) correlations are represented as connecting lines, and only correlations greater than 0.80 are visualized. The overall circos plot representing all correlations in the system is shown in the middle, while simplified circos plots along the top show the correlations with that specific data type. The expression of each biological feature in unstressed (grey) and stressed (black) animals is represented by the outer encircling lines.

Correlation circle plots were employed to visualize the structure of relationships between biological features for each antibiotic treatment condition (i.e., water as a control, Augmentin, and broad spectrum). Variables aligning closely near the outer circle are strongly correlated, while those closer to the center exhibit weaker correlations. Figure 6A illustrates these results for the water group. Many notable taxa clustered tightly next to the outer circle along with certain SCFAs and microglia cytokines, reflecting strong correlations with each other. This included fibre-fermenting taxa (e.g. *Lachnospiraceae*, *Intestinimonas, Anaerotruncus, Bifidobacterium*, *Acetatifactor, Herbinix, Tyzzerella, Lachnoclostridium*, *Oscillibacter*, *Anaerotruncus, Harryflintia,* and *Rosburia*) and SCFAs like propionate, acetate, and butyrate. Strong correlation patterns were also observed between these SCFAs, some splenocyte cytokines (IL-6, IL-10, IL-12 and IL-13), plasma corticosterone, and several microglia cytokines such as, IL-17A/F, IL-15, IL-33, TNF-a, IL-1b, macrophage inflammatory proteins (MIP)-1a and MIP-2, monocyte chemoattractant protein (MCP)-1, IL-9, IP-10, IL-27p28/IL-30, and keratinocyte chemoattractant/growth-regulated oncogene (KC/GRO, aka Chemokine (C-X-C motif) Ligand 1 (CXCL1)), a chemokine responsible for trafficking immune cells and disrupting the blood brain barrier during neuroinflammation.^39^ Thus, these relationships between the microbiome, its production of SCFAs, and microglia, appear highly integrated and may help to coordinate healthy brain development during early life.

When mice were treated with Augmentin, we observed a drastic shift in the correlational relationships between these biological features (Figure 6B). For example, SCFAs and the microbial taxa were no longer correlated to microglial cytokines; instead, these cytokines correlated with each other and plasma corticosterone, apart from other biological systems. Furthermore, hair corticosterone became more closely aligned with colon and some splenocyte cells and cytokines, while many microbiome features shifted closer to the centre of the plot indicating poorer correlations with each other and other biological systems.

Similar to Augmentin, considerable shifts were observed in the interactions between biological systems with broad spectrum antibiotic exposure (Figure 6C). For instance, any interactions with SCFAs are completely absent, consistent with our detected depletion of these important metabolites. Instead, microglial cytokines are strongly correlated with several splenocyte cells and cytokines (e.g. CD25^+^FoxP3^+^Tregs, CD19^+^B cells, CD4^+^ T cells, and IL-10, IL-12, IL-13 cytokines from CD8^+^IL-12, CD4^+^ cells) and body hair corticosterone. Furthermore, microbial clustered strongly with colon cytokines, plasma corticosterone, and select splenocyte cells and cytokines (e.g., IAIE^+^ CD11C^+^ macrophages/dendritic cells, CD8^+^ IFN-g^+^, IL-4^+^). Altogether, these findings suggest that early-life antibiotic treatment disrupts the coordinated interactions between the microbiome, its metabolites, and immune signaling, leading to a loss of system-wide integration that may explain our observed impairment of microglial function and immune homeostasis.

To further examine the interconnectivity among the measured systems, we constructed a similarity matrix and visualized cross-correlations using circos plots for the water, Augmentin, and broad spectrum antibiotic groups (Figure 7A-C). This analysis confirmed previous correlations observed in the previous circle correlation plots and further evidenced the biological effect of stress treatment on the systems evaluated. For instance, within the water treatments, the levels of colon cytokines were higher in stressed vs unstressed animals (Figure 7A). In contrast, correlations observed with key SCFAs (acetate, propionate, butyrate) were more pronounced in water-unstressed (vs stressed) animals. In mice treated with antibiotics (Augmentin or broad spectrum), many of these pairwise correlations were altered compared to the water treatment group (Figure 7B **-C**). For example, fewer pairwise comparisons were observed between individual microbes and other biological features in both antibiotic groups. Among those, much fewer correlations were observed in the Augmentin group, and those detected in the broad spectrum group were mostly negative correlations, suggesting that antibiotic treatment broadly disrupts microbiome-host interactions, leading to weakened system-wide correlations and impaired biological integration.

Overall, this integrative analysis unveils intricate, multi-system interactions among the microbiome, immune, and stress responses. Microbial taxa, SCFAs, and immune cells exhibit strong inter-correlations, underscoring the complex biological network influenced by disruptions to early-life microbial colonization.

## Discussion

This study underscores the profound impact of early-life antibiotic exposure on the gut microbiome and physiological development. Exposure to antibiotics during early life, particularly broad-spectrum antibiotics, significantly disrupts the gut microbiome, drastically reducing microbial diversity and key fermentative microbial taxa, such as *Lactobacillus*, *Akkermansia*, *Lachnospiraceae*, *Bacteroides,* and *Roseburia.* These changes led to reduced SCFA concentrations that persisted into adulthood, revealing the long-term consequences of early microbiome disruption on immune and neurodevelopmental functions. The severe depletion of these microbes and their metabolites from exposure to antibiotics in early-life resulted in altered immune and host responses, most notably an increase in microglia activation in the brain and reduced corticosterone levels. Together, these changes impaired the immune system’s capacity to respond to microbial stimuli and HPA axis function. Antibiotics, rather than stress, were the primary drivers of these disruptions, with some immune alterations persisting into adulthood, thereby increasing susceptibility to allergic airway inflammation. These findings emphasize the critical importance of preserving the early-life microbiome to safeguard long-term immune resilience and physiological homeostasis.

Broad spectrum antibiotics nearly eliminated the gut microbiota, leading to notable reductions in essential commensal bacteria involved in SCFA production. Even when the microbiome showed some recovery by adulthood, SCFAs remained significantly reduced, demonstrating the prolonged impact of early antibiotic exposure on fermentative metabolism. These findings are consistent with prior studies indicating that antibiotics can severely impair the gut’s ability to produce SCFAs, which play vital roles in regulating gut barrier function, immune responses, and brain development.^1,2,40,41^ The persistent reduction of SCFAs despite partial microbiome recovery emphasizes a potential window of vulnerability in early life when the gut microbiota is crucial for shaping long-term host health.

The impact of early-life chronic stress, although less pronounced than antibiotics, was still evident through decreased abundance of key SCFA-producing taxa (*Bifidobacterium* and *Tyzzerella*) and reductions in specific SCFAs, including butyrate, acetate, valerate, and propionate in water-treated stressed mice at P21. This suggests that stress alone can alter the microbiome’s metabolic output, even if overall microbial community shifts are less pronounced. In a related study, prebiotic fibre treatment with 10% oligofructose (which can be fermented to SCFAs) during co-exposure to early-life antibiotics, reversed impaired hypothalamic microglial overactivity in neonatal mice,^40^ further highlighting the importance of SCFA levels for early-life microglial health and supporting neurodevelopment. The reductions in these key metabolites underscore the contribution of stress to altered gut-brain communication and immune modulation during critical developmental periods. However, while stress may exert measurable effects on the microbiome and its metabolites, its impact is overshadowed by the stronger and longer-lasting disruptions caused by antibiotic treatment, reinforcing the predominant influence of microbial depletion on developmental outcomes.

Our modified maternal separation protocol, which started on P7 rather than birth, and gradually increased separation duration, may have mitigated the impact of stress on the microbiome. This approach was designed to reduce the risk of pup cannibalization, as disturbing the litter in the first 3-4 days after birth increases cannibalization in our experience. These modifications may have contributed to the less pronounced effects of chronic stress on the microbiota and physiological state of newborn pups. These findings suggest that the impact of stress on the microbiome is less direct and may require more pronounced or prolonged stress exposure to manifest in measurable changes in developing neonatal pups.

Despite less overt microbiota disruption, stress significantly influenced physiological outcomes, as evidenced by dampened basal corticosteroid levels in plasma and the increased levels in hair at P21, with stress having an additive effect on the antibiotic treatments. These findings are consistent with classical hyperactivation of the stress axis system and negative feedback mechanisms that downregulate circulating stress hormones under chronic conditions. The accumulation of corticosterone in hair suggests that substantial stress exposure was experienced by these pups from birth given that mice are born hairless, and supporting the notion that chronic early-life stress can lead to long-term alterations in HPA axis regulation and stress responsiveness later in life.^42^ Previous studies have demonstrated that the absence of a microbiome leads to dysregulation of HPA axis function, including upregulated expression of corticotrophin-releasing hormone messenger ribonucleic acid (mRNA) in the hypothalamus, reduced glucocorticoid receptor mRNA expression in the hippocampus, and inhibited maturation of hypothalamic microglia, ultimately affecting stress reactivity and neurodevelopmental programming.^43−46^ Our findings, together with other similar studies, collectively highlight the role of the microbiome in moderating stress responses and neurodevelopmental outcomes resulting from changes in its metabolomic output that were not necessarily proportional to overall changes in community structure, further underscoring the complex relationship between microbiome structure and function.

The bi-directional communication between the gut microbiome and the HPA axis play a crucial role in neuroimmune development. Microbial metabolites like SCFAs interact with pattern recognition receptors in microglia and astrocytes, like the nucleotide-binding oligomerization domain-containing protein 2 (NOD2) and peptidoglycan recognition protein 2 (pglyrp2), influencing neurodevelopmental processes.^47^ The increased proportions of the activated, ameboid, and phagocytic microglia in Augmentin and stress treated mice at P21, suggest that these early-life exposures also affect microglial maturation, and consequently, neurodevelopment. This aligns with studies showing that germ-free mice, or those with impaired microbiome composition, exhibit increased immature and activated microglia phenotypes, which are linked to altered brain immune functions and disrupted neural circuit formation during early life.^16,20,48^

Notable changes in microglia cytokine secretion following an LPS challenge in antibiotic- and stressed-exposed mice, highlight the critical role of early-life microbial exposure in shaping microglial function. These findings suggest that early microbiome disruptions impair microglial functions necessary for maintaining central nervous system immune homeostasis. Similar globally-defective microglial immune responses have been reported by Erny *et al*.,^16^ where germ-free mice exhibited impaired microglial function that was rescued using SCFA supplementation. Additionally, Cho *et al*.,^40^ demonstrated that maternal antibiotic exposure during pregnancy and lactation led to offspring with impaired hypothalamic microglial response to LPS. These combined early-life exposures that impair microglial responses to pathogens could have lasting implications for neurodevelopment and brain immune functions later in life.^12,13,16^ Importantly, changes to the immune functions of microglia were more prominent than those detected in spleen or colon, suggesting that microglia are particularly sensitive to early-life microbiome and stress disturbances than other immune compartments.

The observed increase in allergic airway inflammation in antibiotic-treated mice further demonstrates the broader systemic impacts of early-life microbiome disruption from antibiotic exposure. Elevated inflammatory cells in the lungs of Augmentin-treated adult mice following OVA challenge suggest that early-life antibiotic exposure can predispose individuals to heightened inflammatory responses. The long-lasting effects underlying increased allergic airway susceptibility may be mediated by the early-life dysregulation of corticosteroid metabolism, decreased tolerogenic intestinal immune cell populations (i.e., dendritic cells) and impaired microglial function detected in the antibiotic-treated mice, which may have collectively resulted from microbiome disruptions observed during early-life development, or through other mechanisms not measured in our study.

Correlational analysis underscored the systems-level relationships between specific gut microbes, SCFAs, and immune cells populations, further suggesting that SCFAs are pivotal mediators of microbiome-host interactions. Under water treatment conditions, strong positive correlations were noted between acetate, butyrate, and propionate, and many commonly known SCFA-producing microbes like *Bifidobacterium, Intestinimonas, Lachnospiraceae*, *Oscillibacter, Roseburia*, and *Lachnoclostridium*, as well as other lesser known SCFA-producers like *Anaerotruncus*, *Tyzzerella*, *Herbinix, Acetatifactor*, and *Harryflintia*. Interestingly, SCFAs also positively correlated with several LPS-challenged microglial cytokines that are involved with inflammatory propagation, like IL5, IL15, IL-33, TNF-a, IL-1b, MIP-1a, MIP-2, MCP-1, KC/GRO and IL17A/F, with many of these associated with corticosteroid-resistant neutrophilic asthma, lung inflammation, and bacterial infections.^49,50^ SCFAs were also positively and negatively correlated with plasma and whisker corticosterone, respectively, likely a result of opposite corticosteroid response observed in these two tissues, highlighting the complex interplay between microbial metabolites and host endocrine responses that underpin gut-brain axis communication pathways.^2^ Collectively, this suggests a potential connection between peripheral and central immunity that is mediated by microbial production of SCFAs in the gut that dictates early-life development.

However, when these animals were treated with broad spectrum antibiotics, we also observed a drastic shift in the pattern and types of correlational relationships in these interconnected biological systems, whereby, the positive correlations observed between carbohydrate-fermenting taxa, SCFAs, and microglia cytokines (i.e., the gut-brain axis) are lost, likely from the absence of SCFAs. Instead, microbial taxa associated mainly with colon cytokines, potentially indicative of a pro-inflammatory state.

Overall, our data reflect significant interrelationships between the intestinal microbiome, its fermentative by-products, and host immune responses across multiple tissues including the brain. These findings suggest a tightly interconnected system that influences the maturation and function of critical biological pathways during early life. Future research should focus on elucidating the specific molecular mechanisms underlying the observed interactions between the gut microbiome, SCFAs, and immune responses, and connecting changes to the gut microbiome resulting from early-life disturbances to microglia maturity and proportionality through direct experimentation and manipulation of neonatal microglia themselves; a limitation of this study. Identifying specific signaling pathways and microbial metabolites responsible for immune modulation could provide insights into therapeutic targets for restoring the microbiome following antibiotic therapy.

## List of Abbreviations

ASV: Amplicon sequence variants
BALF: Bronchoalveolar lavage fluid
CD: Cluster of Differentiation
CXCL1: Chemokine (C-X-C motif) Ligand 1
EDTA: Ethylenediaminetetraacetic Acid
ELISA: Enzyme-Linked Immunosorbent Assay
EPM: Elevated plus maze
FDR: False Discovery Rate
FACS: Fluorescence activated cell sorting
FBS: Fetal bovine serum
GF: Germ free
HPA: Hypothalamic-Pituitary-Adrenal axis
HBSS: Hank’s balanced salt solution
HRGC: High Resolution Gas Chromatography
IP: Intraperitoneal
IL: Interleukin
KC/GRO: Keratinocyte chemoattractant/human growth regulated oncogene
LPS: Lipopolysaccharide
LSD: Least square means
MCP-1: Monocyte chemoattractant protein-1
MIP-2: Macrophage inflammatory protein-2
NMDS: Non-metric multidimensional scaling
mRNA: Messenger Ribonucleic Acid
MSD: Meso Scale Discovery
NOD2: Nucleotide-binding oligomerization domain-containing protein 2
3NPH: 3-nitrophenylhydrazine
OVA: Ovalbumin
PBS: Phosphate buffered saline
PCA: Principal Component Analysis
PCR: Polymerase chain reaction
pglyrp2: peptidoglycan recognition protein 2
PLS-DA: Partial Least Squares−Discriminant Analysis
RPMI: Roswell Park Memorial Institute
SCFAs: Short chain fatty acids
TMB: 3,3′,5,5′-Tetramethylbenzidine
TNF-α: Tumor necrosis factor Alpha

## Declarations

## Ethics approval

All mouse husbandry procedures were completed under the approval of the Animal Care Committee protocol AC19-0112 of the University of Calgary.

## Consent to participate

Not Applicable

## Consent for publication

Not Applicable

## Availability of Data and Materials

Raw sequences for the current study are available the National Center for Biotechnology Information (NCBI) Sequence Read Archive (SRA) as of the date of publication with accession numbers TBA. Generated scripts are available in the following GitHub repository (TBA).

## Author’s Contributions

VAO (together with MCA) conceptualized, designed and executed the experiments, analyzed associated data, and principally authored the original manuscript with project supervision and editorial assistance from MCA, DF and KAS, who also provided lab space, analytical tools, materials and reagents, and project funding; CM assisted with project design, experimental execution, organ and tissue harvesting and data collection; MRA assisted with data and statistical analyses, sequencing and bioinformatics, and manuscript editing and review; BH assisted with corticosteroid data collection; FAV performed behavioural assessments, associated data collection; EvTB assisted with experimental execution, collection of lung BALF fluid, and histological prep for cell differential analysis; EMM assisted with experimental execution and splenocyte isolation from harvested tissues; KK assisted with experimental execution and maternal separation stress protocols of animals; HD performed short-chain fatty acid determination in mouse stool; VKP assisted with experimental execution, gut tissue harvesting and processing and manuscript editing and review; JS assisted with experimental execution and collection of blood and hair samples. All authors provided editorial feedback and approved the final manuscript.

## Acknowledgments

The authors thank Martha Hurtatis and Gemma Haguisan from the Clara Christie Centre for Mouse Genomics for breeding and care of the mouse colonies; Candida Grivel and Dr. Vanessa Louise Olivier from the Mouse Half-way House for maintaining the mouse colonies during experimentation; and Dr. Lavenia Ionescu, and Yiping Liu from the Flow Cytometry Core Facility for technical assistance.

